# Self-organized acoustic behavior in bats arises from simple rules

**DOI:** 10.1101/2024.11.18.623889

**Authors:** Kazuma Hase, Tasuku Miyazaki, Seiya Oka, Noriyoshi Senoo, Hiraku Nishimori, Masashi Shiraishi, Ken Yoda, Kohta I Kobayasi, Shizuko Hiryu

## Abstract

Although self-organized behavior has been extensively studied in the movement of group-behaving animals, such as fish or birds, far less attention has been paid to vocal behavior in animal groups, especially in mammals. Here, by testing the vocal response of echolocating bats (*Miniopterus fuliginosus*) to the playback of jamming stimuli in the lab, we discovered a mathematical model that can well describe vocal frequency control by bats in response to jamming stimuli mimicking echolocation sounds emitted in a group of bats. We then extended the model to a group of flying bats and observed frequency-adjusting behavior by which the frequency differences for the group increased overall, similar to a previous observation on group-flying bats. Further, we showed the frequency-adjusting behavior led to less potential misdetections by echolocation in a group. Our findings suggest this frequency-adjusting behavior is a self-organized vocal behavior that mitigates conflicts within a group, one achieved via simple behavioral rules, as in other types of collective animal behavior.

## Introduction

Self-organization is a phenomenon whereby the interaction among individuals generates order at the level of a group. In animals, cooperative group behavior emerges when individuals follow several simple rules. For instance, in a school of fish (*1*) or a flock of birds (*2*, *3*), each individual determines its movement based on its position relative to the others, resulting in a coordinated movement of the group as a whole. Studies of such group behavior have primarily focused on how the movement of individuals are governed depending on their positional relationship to others within the same group. Animals perceiving echoes of self-emitted signals to obtain information of surrounding environment, such as echolocating bats, must not only avoid colliding with other bats but also resolve acoustic interference that arises within the group to avoid acoustic confusion or masking due to the calls emitted by the other members. Echolocating bats perceive the world by comparing self-emitted ultrasounds with the returning echoes (*4*), enabling these mammals to track small flying insects in total darkness (*5*). However, within a group, each bat is exposed to the other bats’ sounds, which can impair their respective echolocation ability (*6*, *7*). Although how bats deal with acoustic interference from conspecifics has been studied, there have been no study to describe the coordinated vocal behavior of group-flying bats in a mathematical model.

It is known that bats can change the acoustic characteristics of their echolocation pulses in the presence of other individuals or jamming stimuli. For example, in the field, *Tadarida brasiliensis* flying in a pair emitted pulses whose terminal frequency (TF) differed (*8*). Similarly, in the laboratory, when flying in a group of four, *Miniopterus fuliginosus* bats shift their TF so that the mean TF differences increase in the group (*9*). Importantly as well, these frequency shifts can reduce the similarities of pulses of bats within a given group (*9*). These types of frequency-shifting behavior are consistent with jamming avoidance responses analogous to jamming avoidance responses in weakly electric fish (*10*).

When multiple bats were flying together, even if only a group consisting of relatively few individuals, each bat in the group would experience an ever-changing acoustic scene created by the other bats’ vocalizations. If the frequency-shifting behavior is a jamming avoidance response, how does each bat in the group decide whether, and how, to alter their respective frequency (and of course other acoustic characteristics in tandem) in response to the complex dynamic acoustic scene? Here, we hypothesize that frequency adjustments across an entire group could be autonomously achieved by simple behavioral rules which each member follows.

Here, we conducted behavioral experiments to investigate how bats modified TFs in the presence of jamming stimuli composed of different TFs, interpulse intervals (IPIs), and sound pressure levels (SPLs). This was because frequency, IPIs, and SPLs are considered acoustic parameters able to characterize the acoustic scene created by a group of echolocating bats in flight. These results showed the bats shifted their TFs depending on the IPIs and SPLs. We also found an upward shift in the TF response to stimuli when the surrounding TF was lower than their own TFs, and vice versa (TF shifting slightly downward or not at all in response to a higher surrounding TF). Then, we modeled a leaky integrator model, which has been used to express various biological phenomena such as membrane potentials of a neuron and decision making in animals, to describe and predict these behavioral observations in the presence of the jamming stimuli. Next, by conducting simulations of bats flying in a group using this model, we successfully replicated the frequency-adjusting behavior observed in a previous study (*9*). Altogether, our results demonstrate that frequency-adjusting behavior in bats flying in a group can be explained by relatively simple behavioral rules. In this model, bats do not rely on a fine temporal and spectral resolution of acoustics because their emitted frequency is determined by the imbalance in the surrounding frequency’s power vis-à-vis their own in response to other bats’ pulses received in relatively large time windows. Our findings suggest that self-organized vocal behavior to mediate conflicts in a group could be achieved by members following simple behavioral rules, as in other types of collective behavior, like bird flocking and fish schooling.

## Results

To understand how bats changed the acoustic characteristics of their emitted echolocation pulses in response to different levels of acoustic jamming, we presented solitary flying bats with jamming sounds that mimicked FM echolocation pulses having different IPIs (Fig. S1). Echolocation pulses emitted by flying bats were recorded by a miniature telemetry microphone attached to each bat’s back (*11*). The jamming stimuli consisted of repetitive FM jamming sounds, whose bandwidth was 40 kHz with 3-ms duration, and the shape of their frequency sweep simulated FM echolocation pulses. We generated two types of jamming stimuli, with low versus high TFs compared to the overall mean TFs of *M. fuliginosus*.

Evidently, bats shifted the TFs of their emitted pulses upward in response to jamming stimuli with a low TF, and these upward shifts increased as the IPIs of jamming stimuli increased (ANOVA: *F*_4, 30_ = 11.6, *P* = 0.835; Fig. 1a, b and Fig. S2; results for other acoustic characteristics can be found in Fig. S5, S6). The shifts were significantly different between shorter and longer IPIs (respectively, 50 ms vs. 200, 800, or 1600 ms and 100 ms vs 800 or 1600 ms; Tukey-Kramer, all *P*-values < 0.05; Fig. 1b). Whereas the mean TF shift was 1.4 ± 0.8 kHz when IPI of the jamming stimuli was the shortest (IPI: 50 ms), it was much lower, at 0.0 ± 0.2 kHz, in the presence of jamming stimuli with the longest IPI (IPI: 1600 ms). Because the IPI of jamming stimuli can represent the density of animals in a group, these results demonstrated that echolocating bats were able to shift their TF according to their group size.

**Figure 1.**
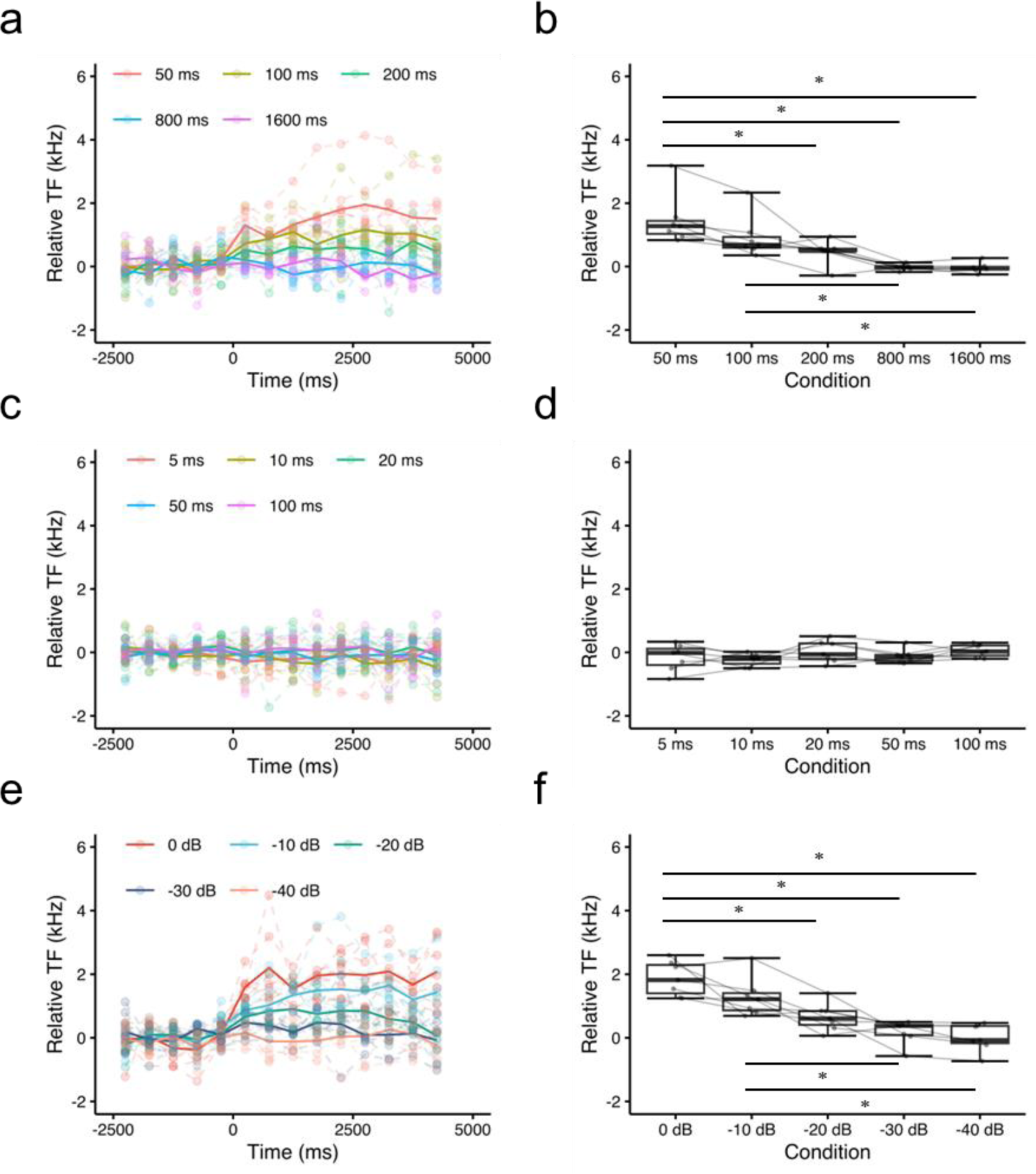
Echolocation behavior of flying bats exposed to jamming stimuli. Changes in the terminal frequency (TF) of emitted pulses by bats in response to low TF-jamming stimuli with different IPIs (**a, b**), high TF-jamming stimuli with different interpulse intervals (IPIs) (**c, d**), low TF-jamming stimuli with different sound pressure levels (SPLs) (**e, f**). For each condition, n = 7 bats. Dashed lines and solid lines indicate individual mean TF and the time course of changes in population mean TF. Colors correspond to different stimulus parameters in **a, c, e**. Plots and lines show mean changes in the TF of each individual in **b, d, f**. For each boxplot (**b, d, f**), the medians are indicated by horizontal lines in the boxes, whose upper and lower bounds respectively show the first and third quartiles, with whiskers bars above and below showing the range of values within a 1.5 interquartile range. Asterisks indicate significant differences detected by the Tukey–Kramer multiple comparison test.

Next, we examined how jamming stimuli with a high TF may have affected the acoustic characteristics of echolocation pulses. With respect to *M. fuliginosus*, our recent study revealed they did not change their TF in response to jamming stimuli that had a TF higher than their own (*12*). By contrast, Hase et al. (2018) observed that some individuals of that species shifted their TF downward when flying in a group (*9*). Since we detected here upward shifts caused by jamming stimuli with a low TF that depended on the IPIs of stimuli, we hypothesized that high TF-jamming stimuli with shorter IPIs could elicit downward shifts in the bats’ TF. To test that, we used high TF-jamming stimuli with shorter IPIs than those used for low TF-jamming stimuli. Overall, the imposed high TF-jamming stimuli failed to induce significant shifts in TF (ANOVA: *F*_4, 30_ = 1.2, *P* = 0.3153; Fig. 1c, d and Fig. S3). The mean TF shift was –0.2 ± 0.4 kHz in response to the shortest IPIs; however, 2 of 7 tested bats exhibited slightly decreased TFs under extreme conditions (please refer to bat 601 and 603 in Fig. S3). This latter result implied bats might change their TFs bi-directionally and asymmetrically according to spectral relationships between their own TF and the TFs of surrounding jamming stimuli. Another factor by which to gauge acoustic jamming is SPLs, so we then exposed bats to jamming stimuli with differing SPLs. In this experiment, only a low TF jamming stimuli was applied, given the above negligible effects of high TF-jamming stimuli on TF. Evidently, those observed upward shifts in the TFs of bats’ emitted pulses decreased as the SPLs decreased (ANOVA: *F*_4, 30_ = 17.64, *P* < 0.05; Fig. 1e, f and Fig. S4). Notably, the shifts differed significantly in magnitude among SPLs (0 dB vs. –20, –30, or –40 dB, and –10 dB vs. –30 or –40 dB; Tukey–Kramer: all *P*-values < 0.05; Fig. 1f). They did not shift TFs in response to jamming stimuli when the stimuli were presented at –40 dB relative to the maximum SPL (mean TF shift: 0.0 ± 0.4 kHz).

The IPIs of jamming stimuli can indicate the number of or the density of individuals in a group, while the SPLs of jamming stimuli can be used to roughly gauge the distance among group members. Also, because every bat in a group receives pulses from other members entailing inter-individual variation, the TFs of jamming stimuli are related to relation of TFs among individuals of that group. Therefore, our behavioral results led us to formulate the following hypothesis. First, the bat with the highest TF in a group is most affected by other individuals because they all have lower TFs than it, so it responds and changes its TF the most; likewise, the bat having the second highest TF is the second-most affected and so its change in TF should be the second greatest in magnitude; conversely, the bat with the lowest TF is least affected, hence its TF is changed the least in magnitude. In this way, as a group, these bats could achieve a frequency-adjusting behavior autonomously, in the absence of any central command (Fig. S7).

To test that hypothesis, we constructed a mathematical model describing changes in TF in response to jamming stimuli, and then applied the model to more realistic situation, that is, group flight. We described the bat responses to jamming stimuli via a leaky integrator model (Fig. S8). This model worked as follows. First, the input was a rectangular pulse train representing overall power of each jamming stimulus. The model then integrated the pulse train with a leak. Next, the TF was determined by multiplying the integration with a scaling factor whose value depended on differences in the TFs between a bat and a jamming stimulus.

Figure 2 shows how the model operates in the presence of various types of jamming stimuli. Overall, the model gave results similar to those observed in the behavioral experiments. The modeled bats shifted their TFs upward in response to low TF-jamming stimuli, but these shifts decreased as the IPIs of jamming stimuli increased (Fig. 2a, b). The modeled bats did not modify their TFs in response to high TF-jamming stimuli, though we discerned slight downward shifts under the shortest IPI condition (Fig. 2c, d). Importantly, the upward shifts in the TFs of modeled bats in response to low-TF jamming stimuli decreased as the SPLs decreased (Fig. 2e, f). These data confirmed the model could capture fundamental features of behavioral responses by real bats to simulated acoustic interferences mimicking a group flights.

**Figure 2.**
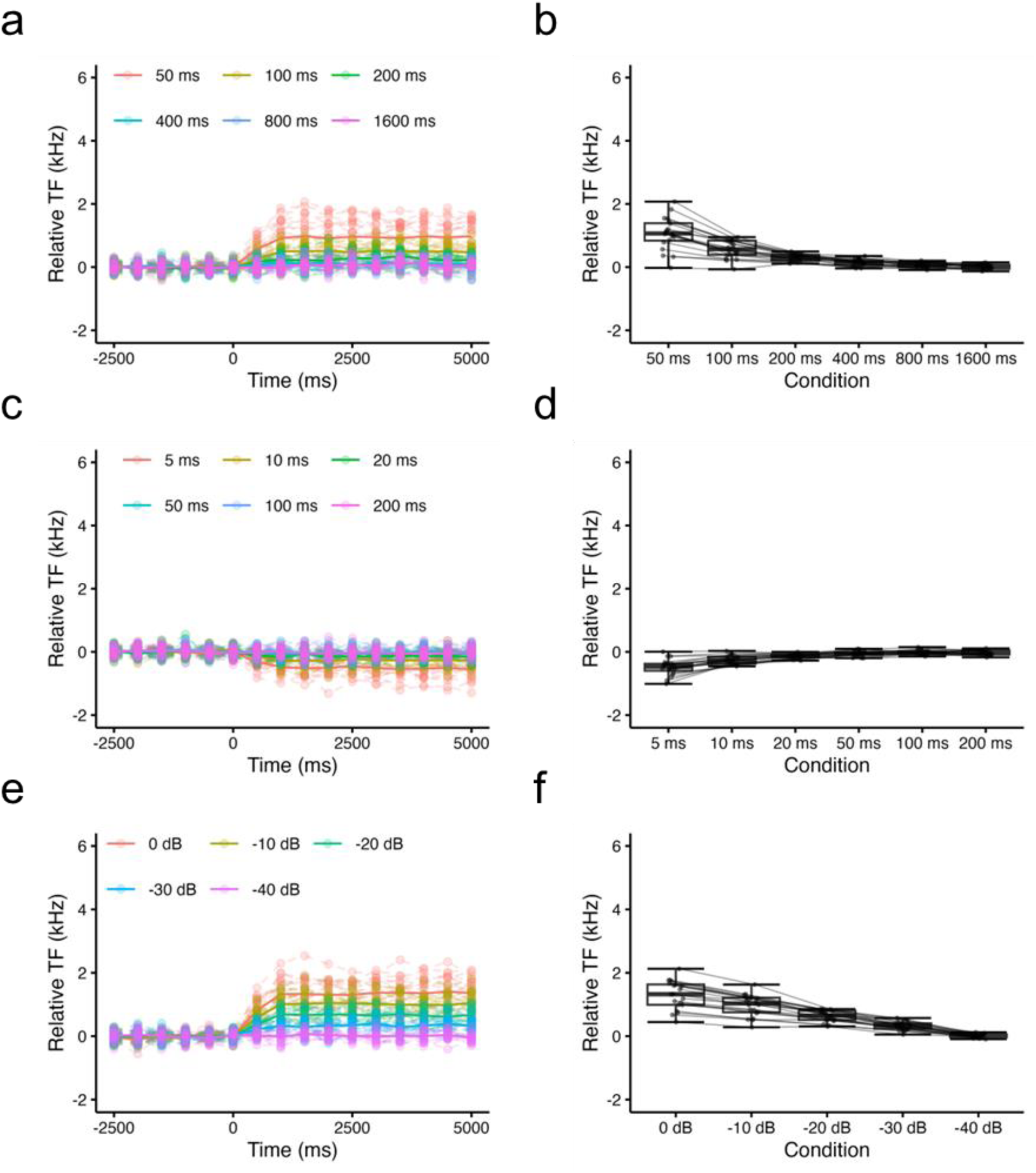
Modeling the behavior of flying bats in the presence of jamming stimuli. Changes in TF reproduced by the model in response to low TF-jamming stimuli with different IPIs (**a, b**), high TF-jamming stimuli with different IPIs (**c, d**), low TF-jamming stimuli with different SPLs (**e, f**). For each condition, n = 20 model evaluations. Dashed lines and solid lines indicate individual mean TF and the time course of changes in population mean TF. Colors correspond to different stimulus parameters in **a, c, e**. Plots and lines show changes in the mean TF of each evaluation in **b, d, f**.

Then, the model was extended to simulate the behavior by group-flying bats. We first generated flight trajectories of four bats in a 5 m × 5 m space to simulate the effect of the atmospheric attenuation and directionality of emitted pulses. Each individual had its own TF as an initial value, generated from a normal distribution. The modeled bats emitted pulses with IPIs, the latter determined by a gamma distribution (Fig. S9a). For the sake of simplicity, the bats emitted pulses at a constant SPL. We estimated the SPLs of pulses that one bat received from co-occurring bats by considering atmospheric attenuation and directionality of emitted pulses, these calculated based on the distances and angles among individuals (Fig. S9b, c). Figures 3 shows two examples of the simulation applied to a group size consisting of four bats (Fig. 3a, b). For both, the modeled bats’ differences in their TFs became greater going from solo to group flight (Fig. 3c, d). We repeated the simulation 100 times, finding that the individual differences in mean TFs rose from 0.6 ± 0.6 [95% Confidence interval (CI): 0.54–0.72] kHz and 0.7 ± 0.6 [95% CI: 0.57–0.75] kHz in single flights 1 and 2, respectively, to 1.3 ± 0.7 [95% CI: 1.18–1.41] kHz in the group flight (Fig. 3e). This pattern is similar to what we reported in a previous behavioral study, in which groups of four bats had increased individual differences in their mean TFs in group flight (*9*). The same simulations were performed with a group size of 8 (Fig. 4), 16, and 32 bats. This revealed a similar pattern in the larger groups. The frequency differences among individuals were 0.3 ± 0.3 [95% CI: 0.34–0.41], 0.2 ± 0.2 [95% CI: 0.20–0.23], and 0.1 ± 0.2 [95% CI: 0.12–0.14] kHz in single flight 1, but increased to 0.9 ± 0.6 [95% CI: 0.79–0.92], 0.5 ± 0.5 [95% CI: 0.45–0.52], and 0.3 ± 0.4 [95% CI: 0.26–0.30] kHz in the group flight for a group size of 8, 16, and 32, respectively.

**Figure 3.**
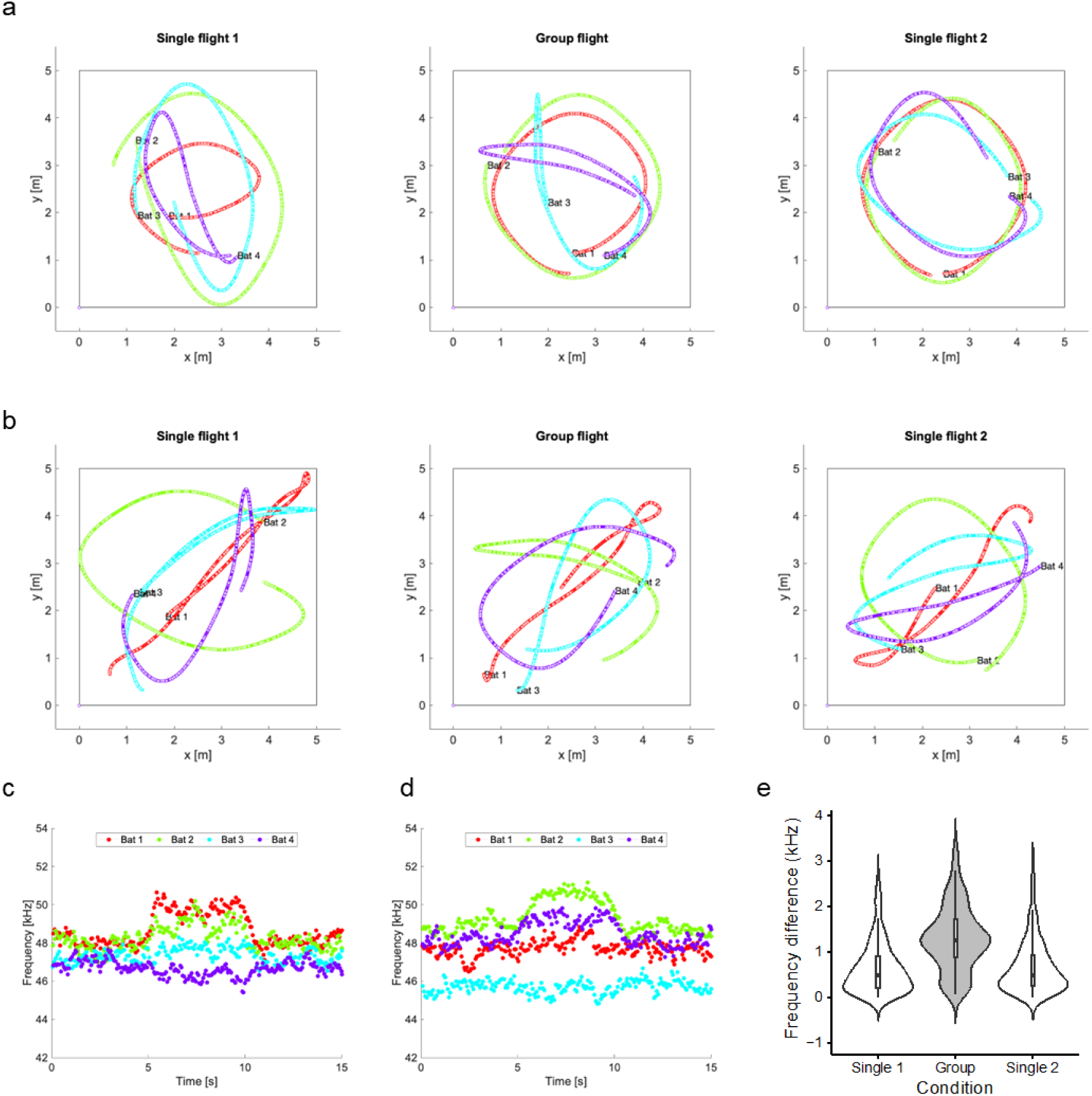
Modeling the behavior of flying bats in a group size of four bats. Two examples of the simulations of four bats flying in a group in a two-dimensional 5 m × 5 m space. In (**a, b**) are generated flight trajectories of four bats in example 1 (**a**) and 2 (**b**). The bats are flying together yet are unaffected by each other in the first 5 sec (‘Single flight 1’; left column in **a, b**); then they are affected by each other for 5 seconds (‘Group flight’; middle column); after that, they are again flying unaffected by the other bats (‘Single flight 2’; right column). Solid lines with different colors represent the flight trajectory of different individuals in each group. Plots on the lines indicate when the bats emitted pulses. **c, d,** Changes to TFs in the simulations in the examples 1 (**c**) and 2 (**d**). In both simulations, bats emitted pulses with slightly different TFs in both Single flights and augmented their inter-individual differences in mean TFs during the Group flight. Panel (**e)** depicts the changes to the inter-individual differences in mean TFs that resulted from 100 simulations in total.

**Figure 4.**
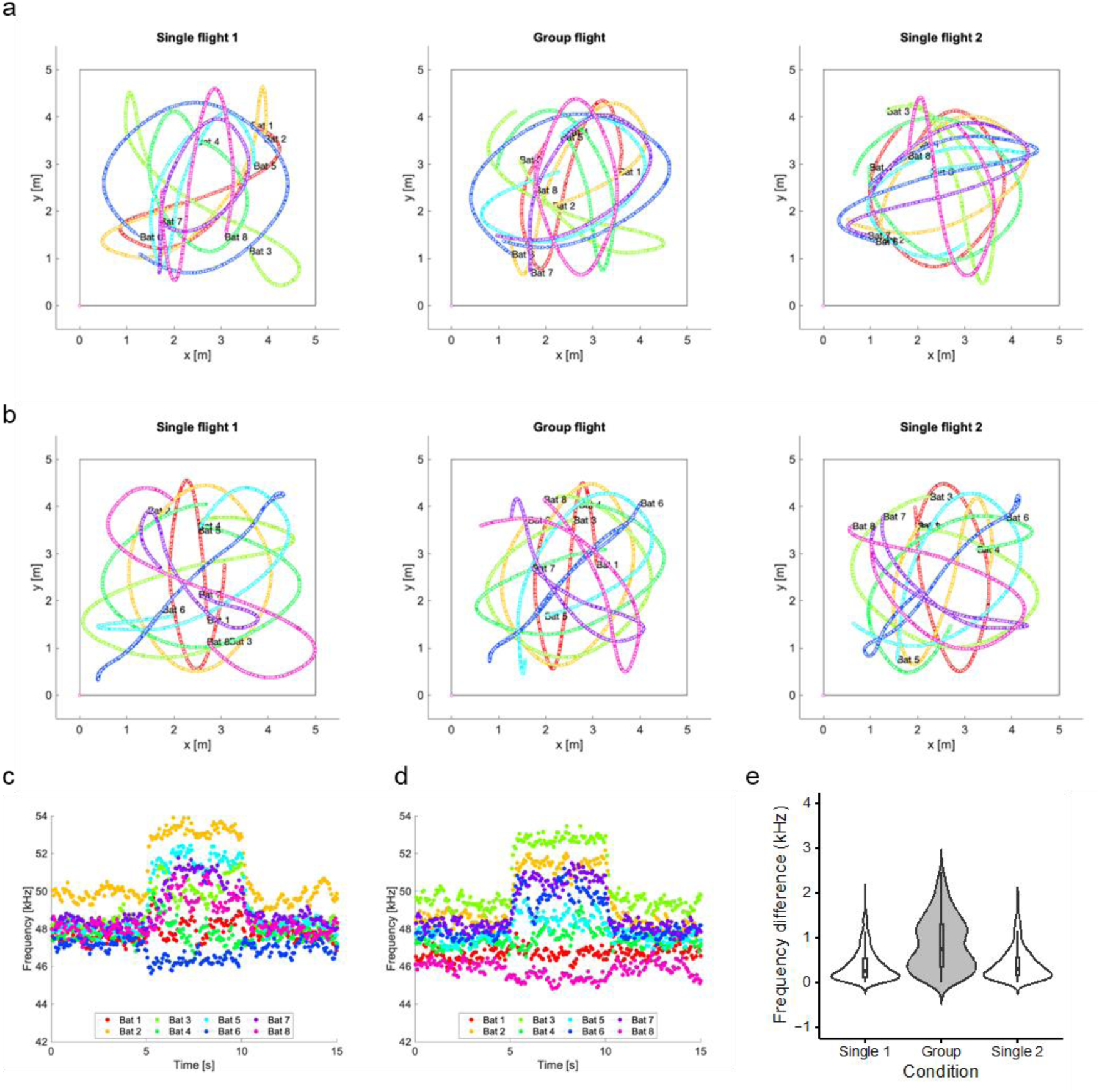
Modeling the behavior of flying bats in a group size of eight bats. Two examples of the simulations of eight bats flying in a group in a two-dimensional 5 m × 5 m space. In (**a, b**) are the generated flight trajectories of four bats in example 1 (**a**) and 2 (**b**). The bats are flying together yet are unaffected by each other in the first 5 sec (‘Single flight 1’; left column in **a, b**); then they are affected by each other for 5 seconds (‘Group flight’; middle column); after that, they are again flying unaffected by the other bats (‘Single flight 2’; right column). Solid lines with different colors represent the flight trajectory of different individuals in each group. Plots on the lines indicate when the bats emitted pulses. **c, d,** Changes to TFs in the simulations in the examples 1 (**c**) and 2 (**d**). In both simulations, bats emitted pulses with slightly different TFs in both Single flights and augmented their inter-individual differences in mean TFs during the Group flight. Panel (**e)** depicts the changes to the inter-individual differences in mean TFs that resulted from 100 simulations in total.

Lastly, to estimate the effect of jamming avoidance responses (JARs) on echolocation performance, we tallied the number of possible misdetections that occurred in the above simulations of group-flying bats (Fig. 5). First, we calculated cross-correlation function between each pulse and other bats sounds detected in a given time window (each spanning the period from current pulse emissions to the next pulse emissions). For this estimation, we normalized all the sounds so they had the same amplitude, rendering the cross-correlation values to range from 0 to 1. The modeled bats were presumed to detect pulses when the peaks of cross-correlation values exceeded 0.5 (Fig. S10). The frequency differences among group members decreased as the group size increased, and they were larger in the simulated groups where each individual employed JARs. The number of possible misdetections fell from 0.14 ± 0.14 to 0.01 ± 0.03 for the 4-bat group, from 1.37 ± 1.07 to 0.12 ± 0.12 for the 8-bat group, from 5.98 ± 2.67 to 2.49 ± 2.33 for the 16-bat group, and from 15.77 ± 4.34 to 10.01 ± 6.11 for the 32-bat group, when JARs are employed by all members (Fig. 5f).

**Figure 5.**
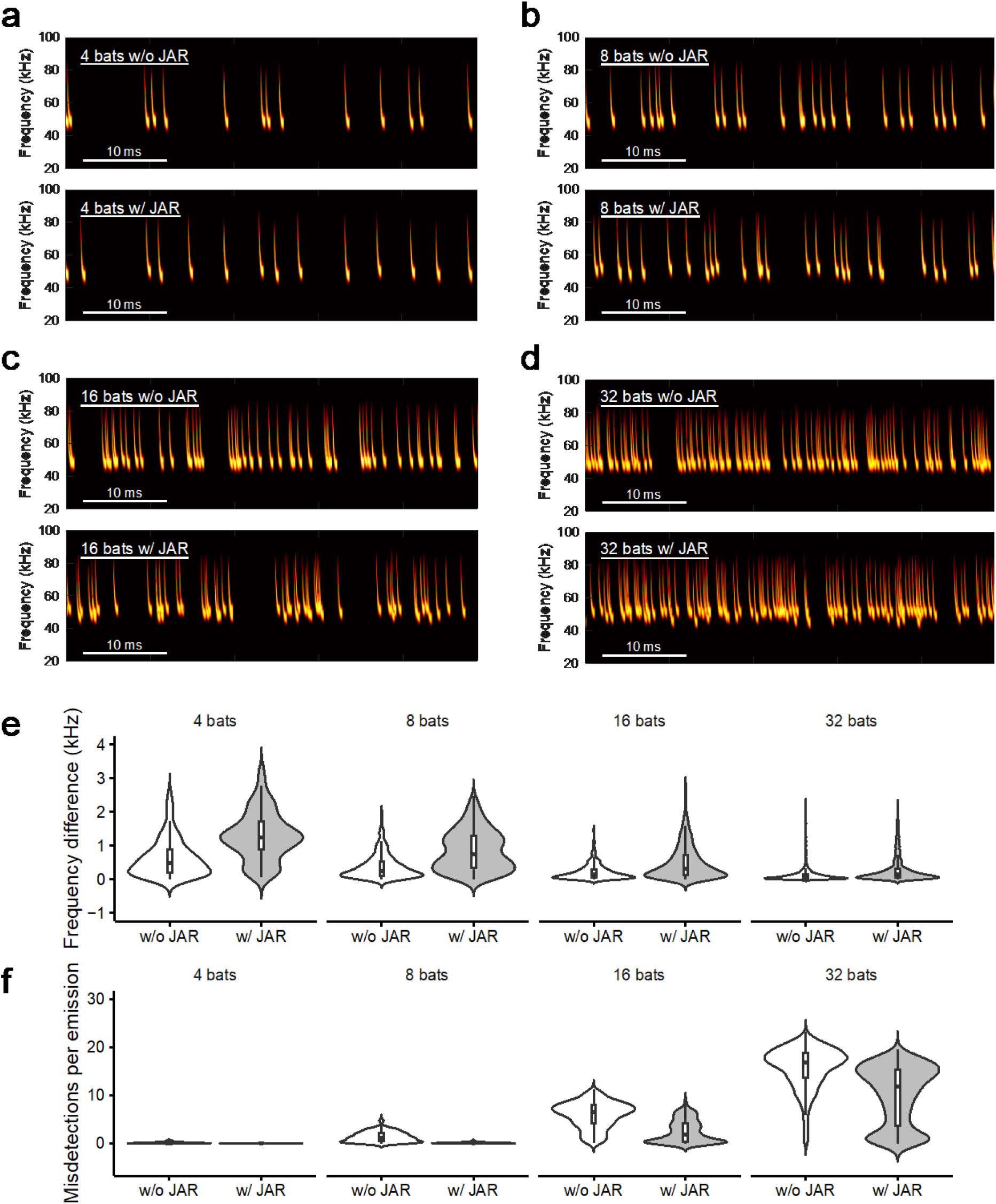
Relationship between bats’ group size and frequency-shifting behavior. Examples of spectrograms of pulses a bat receives from the other group members in each simulation for a group size of four (a), eight (b), 16 (c), or 32 (d) bats. Panel (e) shows the frequency differences among individuals in a group, with and without JAR (Jamming avoidance response). In (f) are the tallied possible misdetections in each group flight, with and without JAR. Violin plots indicate kernel densities of each variable for a different group size and the presence/absence of JAR.

## Discussion

Animals need to modify their behavior to avoid conflicts within a group, often doing so by modulating signals so these can be transmitted effectively among conspecifics. For instance, male frogs alter their vocal timing in a dense mating aggregation (*13*, *14*), and fireflies synchronize the timing of their flashing when they reach a high density (*15*). Those are well-known examples of self-organized signaling in aggregated animals. Bats are social mammals and often behave in a large group; yet how they echolocate while grouped is still mostly unknown because tracking each individual is challenging. Here, by using echolocating bats as a study species, we examined how their self-organized vocal behavior enables them to avoid acoustic interference that could arise during group flight (*7*, *16*). Through acoustic measurements of flying bats recorded by a miniature telemetry microphone, coupled with a mathematical model built based on those results, we obtained the following observations and implications. First, bats increase the upward shifts in their TF as the IPI of the jamming stimuli decreases and the SPL increases. Second, these shifts depend on the frequency of the jamming stimuli. Third, the constructed leaky-integrator model describes well the basic behavioral features of bat responses to the jamming stimuli used in our study. Fourth, this model could also be applied to multi-bat situations and robustly reconstructed results from a previous study in which four *M. fuliginosus* flew together in the lab (*9*). Our model simulations show frequency-shifting behavior increases the inter-individual differences in the TF and lessens the possible number of misdetections by echolocation as a group. Our results are the first to describe how frequency-adjusting behavior in group-flying bats can be a self-organized vocal behavior based upon simple rules.

Leaky-integrator models have been utilized to study decision-making in noisy environments by vertebrates, such as zebrafish (*17*) and primates (*18*), and vocal control in echolocation by bats (*19*). A leaky-integrator model can explain well the involuntary increases in the amplitude of pulses emitted by resting *Eptesicus fuscus* in response to broadband noise (i.e., the Lombard effect), demonstrating that the amplitude can be adjusted within 60 ms from the start of the noise presentation. These findings suggest that the Lombard effect and frequency-shifting behavior share similar mechanisms of audio– vocal integration involving processes achieved by a leaky-integrator.

Horseshoe bats (*Rhinolophus ferrumequinum*) compensate for Doppler shifts in echo frequency by lowering their pulse frequency to maintain an echo frequency within a narrow frequency range, called the reference frequency, where their auditory sensitivity and frequency resolution are extremely high (*11*, *20*, *21*). Over 20 years ago, Metzner et al. (2002) introduced a conceptual audio-vocal feedback model to explain how bats achieve the Doppler shift compensation (*22*). We propose a similar model wherein the pulse frequency is determined by integrating the power of a jamming stimulus with some leak in auditory nuclei with the inputs (or outputs) to the nuclei scaled by the frequency of the stimuli. The model simply relies on the spectral information and does not require extraordinary spectro-temporal resolutions in the auditory system. Also, the model realizes self-generated frequency adjustments without higher-order processes, such as decision-making, to mitigate acoustic jamming in a group of flying bats.

We found that two out of seven tested bats showed downward shifts in their TFs in response to high TF-jamming stimuli with shorter IPIs. Several studies have shown that bats shift their TF downward in response to jamming stimuli (*23*, *24*), and our own previous work also suggests that some individuals decrease their TF in a group of four bats (*9*). Here we observed notable downward shifts in TF in the simulation, likely because we included asymmetrical effects from other bat pulses having a lower TF and those having a higher TF (see Fig. 3c, d and Fig. 4c, d) as in a real group of flying bats. Because downward shifts are uncommon, we suggest that echolocating bats are affected by the presence of pulses whose TFs are higher than their own TFs, and the resulting effect can sometimes be discerned as downward shifts in TF when that effect is strong enough. Further detailed investigation is needed to understand how jamming stimuli with higher TF affects the behavior of bats and also their internal state.

Several studies have suggested that frequency shifts observed in response to acoustic interferences from speakers or conspecifics could be a by-product of different mechanisms. Animals including frogs (*25*), birds (*26*), primates (*27*), and bats (*12*, *19*, *28–30*) display the Lombard effect, which occasionally entails changes in other characteristics such as an increase in duration and/or frequency of vocalization. It is suggested that the frequency shifts we observed in bats could be accounted for by the Lombard effect (*31*). Other studies argued that such frequency shifts are a by-product of attention to nearby conspecifics (*32*). As they approach an object, echolocating bats decrease the pulse duration (*11*, *33–35*), and the pulse duration and the TF are negatively correlated (*32*, *35*). Therefore, if a bat pays attention to nearby conspecifics, the pulse duration become shorter, increasing the TF. The present study does not exclude those hypotheses. Regardless of the mechanisms involved, our study nonetheless provides a novel integrated sensory-motor model that can explain the frequency shifts observed in many previous studies. As long as bats change the TF in the presence of conspecifics, by whatever mechanisms, inter-individual variation in the TF can become greater, which could lead to similar results as reported here (Fig. 5), which ought to be generally helpful for segregating signals from those of conspecifics in groups.

What is the possible functional role of TF shifts? One of the simplest hypotheses is that bats diverge in emitted frequencies to avoid spectral overlapping of their own echoes with other sounds from co-occurring bats. Indeed, our simulation reveals that the number of misdetections is substantially reduced during the frequency-shifting behavior (Fig. 5). Ultrasonic pulses transmitted from a bat to other conspecifics are attenuated in a way that depends on differences in the distances and angles between these bats (*36–38*). Moreover, bats might process only sounds occurring within a short time window after they emit a pulse (*39*, *40*). In addition, inter-individual differences in pulse characteristics can also influence the similarity of sounds among members in a group (*31*, *41*). Furthermore, bats are known to alter the duration (*9*, *31*, *42*), amplitude (*9*, *12*, *31*) and interpulse interval of pulses in response to sounds of conspecifics (*43–46*). Thus, those passive and active variations in pulse acoustic characteristics will lessen the number of misdetections that can be further reduced with the self-organized frequency-adjusting behavior, enabling bats to echolocate in a large group.

## Materials and Methods

### Subjects

A total of 11 *Miniopterus fuliginosus* (six males and five females) were used in this study. They were caught from wild colonies in caves in Fukui Prefecture, Japan. We were licensed to collect these bats, carried out in compliance with all Japanese laws (permits from Fukui Prefecture in 2020, 2021, and 2022). The bats were housed in a temperature- and humidity-controlled room [4 m (L) × 3 m (W) × 2 m (H)] at Doshisha University, Kyoto, Japan. Food (meal worms) and water were made available *ad libitum* to the bats. The day/night cycle of the room was set to 12-h/12-h. All experiments were conducted in accordance with all Japanese laws. All experimental protocols were approved by the Animal Experiment Committee of Doshisha University.

### Experimental design

The experimental procedures used followed those applied in several previous studies (*12*, *42*, *47*). Recordings were made in an experimental chamber (9 m × 4.5 m × 2.4 m) at Doshisha University, Kyoto, Japan. This experimental chamber was constructed with steel plates to reduce external electromagnetic noise and signals from commercial FM radio stations. Its inside walls were lined with sound-attenuating foam, to reduce acoustic reflections. A telemetry microphone (*11*, *48*) was attached to the back of a bat to record its emitted pulses in flight (for the details, please refer to the following section). During the experiments, long-wavelength lighting with filters (removing wavelengths < 650 nm) was used to prevent a bat from using visual cues (*49*). Solitary bats were flown in a flight space (3.0 m × 4.5 m × 2.4 m) delimited by a net suspended from the ceiling. Four loudspeakers (Pioneer Corp., PT-R7 III, Kanagawa, Japan) were set at a fixed height (1.4 m) in each corner of the flight space (within the chamber).

First, we flew a bat in the flight space for ca. 20 sec, and then exposed it to a jamming stimulus (5 sec) while flying; after the presentation of the jamming stimulus ended, the bat kept flying for another 20 sec. During this procedure, none of the 11 bats landed on the walls, the ceiling, or the floor, though some bats did occasionally try to land somewhere in the flight space. We considered the 5-sec period before the presentation as the “jamming off” condition, and the 5-sec period of the presentation of jamming stimulus as the “jamming on” condition. We tested 7 bats once for each of the three different stimuli.

### Sound recordings

Echolocation pulses emitted by a bat during flight were recorded with a miniature telemetry microphone (*48*). It consisted of a 1/8-inch omnidirectional electret condenser microphone (Knowles, Model FG-23329-C05, Itasca, IL, USA) and a custom-made amplifier and transmitter circuit with a transmitting antenna, powered by a 1.5-V hearing aid battery (Sony, Type SR626SW or SR726SW, Tokyo, Japan). Including the battery the telemetry microphone weighed ca. 0.7–0.9 g, this being just 3% to 7% of a bat’s weight, and thus easily carried when it flew (*9*). Just prior its flying trial, to the back of each bat, we gently attached the telemetry microphone with a piece of double-sided adhesive tape; then, after completing each experiment, we carefully removed the microphone with a parting agent (Toagosei Co., Ltd., Aron Alpha Specific Peeling Liquid, Tokyo, Japan).

The telemetry microphone transmitted FM radio signals having a carrier frequency between 76 and 104 MHz, depending on a resonant frequency of the circuit. These signals were received with an FM radio antenna (Terk Technologies Corporation, FM+, Commack, New York, USA) suspended from the ceiling and demodulated and then band-pass filtered (10–200 kHz) with a custom-made multi-channel FM receiver (ArumoTech Corporation, Kyoto, Japan). The de-modulated signals were then digitized with a USB data acquisition device (16-bit, fs = 500 kHz; Model NI USB-6346 or Model NI USB-6356, National Instruments, Austin, TX, USA).

### Jamming stimuli

Jamming stimuli consisted of sequences of downward FM signals differing in terminal frequency (TF), sound pressure levels (SPLs), or interpulse intervals (IPIs). These FM signals, designed to mimic bats’ echolocation pulses according to previous studies (*47*, *50*), were 3 ms in duration and 40 kHz in bandwidth. We prepared two types of FM signals in terms of TF. One was modulated from 85 to 45 kHz (‘low TF’) and the other from 95 to 55 kHz (‘high TF’), by using the mean TF of *M. fuliginosus* flying in the chamber (approximately 48 kHz) as a cut-off. To understand how bats respond to different IPIs of the jamming stimuli with contrasting TFs, we generated the low TF stimuli with IPIs of 50, 100, 200, 400, 800, and 1600 ms and the high TF stimuli with IPIs of 5, 10, 20, 50, and 100 ms. A different set of IPIs for low and high TF stimuli was used because we hypothesized the high TF stimuli, which usually did not induce shifts in TF (*12*), would cause shifts in the TFs of pulses emitted by bats if they were presented with higher repetition rates,. The SPLs used here were 0, –10, –20, –30, and –40 dB (relative to the maximum SPL) for both low TF and high TF stimuli with an IPI of 100 ms. The maximum SPL of the jamming FM signals was set at 110 dB SPL, peak-to-peak, at 0.1 m from each loudspeaker.

### Sound analysis

Echolocation sounds emitted by the bats were manually analyzed from sound spectrograms of the telemetry microphone recordings, by using custom-written MATLAB scripts on a personal computer. We defined the TF as the lowest frequency in the spectrograms that was −25 dB from the maximum energy portion of each pulse. The IPIs and their duration were also determined from the spectrograms at −25 dB relative to the maximum energy portion of each pulse.

### Statistical analysis

We used linear mixed models (LMMs) to test whether the TF, pulse duration, or initial frequency (IF) of echolocation pulses were affected by the IPIs or SPLs of the jamming stimuli. We built separate LMMs for three different conditions (i.e., IPI conditions with a low or high TF and the SPL condition with a low TF). The response variables were changes in the TFs, durations, or IFs of sounds emitted by the bats. The fixed effects were the variable (IPIs or SPLs), the presence (jamming off or on) of the jamming stimuli, and the interaction of these two effects. The bat ID was included as a random effect. In the case of significant main effect or interaction, post hoc tests were performed using the Tukey–Kramer method to adjust the Type I error control to correct for the number of multiple comparisons.

The model fit was quantified by plotting residuals. A type II Wald χ2-test was used to detect a significance of the fixed effects. In the case of significant main effect or interaction for either, post hoc tests were performed using the Tukey–Kramer method to adjust the Type I error control to correct for the number of multiple comparisons. All statistical analyses were implemented in the R platform (v. 4.3.0) (*51*). We used the lmer function of the ‘lme4’ package (v. 1.1.33) for building LMMs (*52*), and the Anova function of the ‘car’ package (v. 3.1.2) for the type II Wald χ2-tests (*53*). The model fit was quantified by using the function simulateResiduals of the ‘DHARMa’ package (v. 0.4.6) (*54*). The emmeans function of the ‘emmeans’ package (v. 1.8.6) was used to perform the post hoc comparisons (*55*). *P*-values < 0.05 were considered significant. Results are presented as mean ± SD, unless otherwise stated.

### Modeling the frequency-shifting behavior in response to jamming stimuli

A leaky integrator model was used to simulate the frequency shifting behavior by a flying bat in response to the jamming stimuli, using this equation:

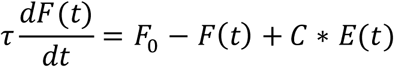

where *τ* is the leak time constant of 200 ms, this empirically determined from results of previous studies that applied jamming stimuli (*12*, *47*, *56*); *F*_0_ is the baseline TF of the bat; *F*(*t*) is the TF of emitted pulses by the bat; *E*(*t*) is the effect of jamming stimuli, and *C* is a scaling factor.

To convey the effects of jamming stimuli, we used rectangular pulse trains (Fig. S8). The IPIs and SPLs of the imposed jamming stimuli were then characterized by deriving the IPIs and amplitudes of those rectangular pulses. Each rectangular pulse lasted 5 ms. We adjusted the scaling factor for low TF and high TF stimuli depending on behavioral experiments’ results obtained here; these *C* values factors were used to model the frequency-shifting behavior in group flight. Random fluctuations in the TF were added to the TF of each bat. We applied the Runge–Kutta 4^th^ order method to solve the above equation, using a 1-ms step. For each stimulus condition, we ran the simulation 20 times.

### Modeling the frequency-shifting behavior in group flight

We generated flight trajectories of bats in each group for 15 sec in a two-dimensional space, by using the following equations:

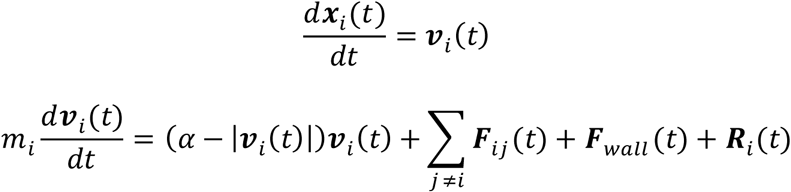

where ***x***_*i*_ is the two-dimensional position of each *i*-th bat; ***v***_*i*_ is the two-dimensional velocity of each *i*-th bat, *m*_*i*_ is a mass of each *i*-th bat, and *α* is self-propulsion. The term ***F***_*ij*_ (*t*) denotes the repulsive forces acting on each bat from other individuals in the group, ***F***_wall_ (*t*) is the repulsive force from the nearest wall, and ***R***_*i*_(*t*) is a random factor. When bats were affected by each other (i.e., *E*_*i*_(*t*) = 0 in the equation bellow) from 5 to 10 sec of the 15-sec flight period, it was designated a group flight. Accordingly, the period before and also after a group flight, when they did not affect each other, were defined here as single flight 1 and 2, respectively. The baseline frequency for each bat in a group was sampled from a normal distribution with a mean of 48 kHz and one standard deviation of 0.4 kHz determined from a previous study (*9*). The IPIs of each bat were sampled from a gamma distribution having a mean of 70 ms and one standard deviation of 18.7 ms (Fig. S9a), this roughly determined from previous studies (*9*, *12*).

The modeled bats were assumed to emit pulses in their flight direction and at a constant SPL. The SPLs received by a bat from other bats during the group flight were calculated by considering attenuation due to the spatial relationships between individuals. The spreading loss and atmospheric attenuations (*36*) were calculated based on the distances between the bats (Fig. S9b, c). We used 50 kHz for calculations of all frequency dependent attenuations and directionalities, since slight differences in frequency negligibly altered the attenuation levels.

We then built a model for frequency-shifting behavior in the group flight of bats, using these equations:

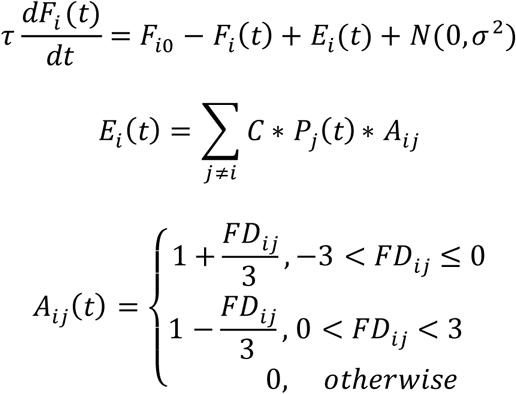

where *τ* is the leak time constant (200 ms); *F*_*i*0_ is the baseline TF of the *i*-th bat; *F*_*i*_ (*t*) is a TF of the *i*-th bat at time *t*; *N*(0, *σ* ^2^) denotes the random changes in the TF, by sampling from a normal distribution; *E*_*i*_(*t*) is a jamming effect from other bats on the *i*-th bat; *P*_*i*_(*t*) is normalized power of pulses received by the *i*-th from other bats, and *C* is a scaling factor, which changes depending on whether the differences between own TF is higher or lower than those of others. In the present study, *C* was set to −4 and 20 when a bat TF was lower or higher than another bat, respectively. The *A*_*ij*_(*t*) term is the factor that determines how the *i*-th bat is affected by *j*-th bat, and *FD*_*ij*_ is the frequency difference between *i*-th and *j*-th bats (Fig. S9). We performed 100 simulations, using a time step of 1 ms, in Matlab 2021b software. We reported 95% confidence intervals (CIs) for the differences in mean TF as well as means ± SDs.

### Estimating the effect of JARs on echolocation in a group

For the simulations of group flight, we estimated possible numbers of misdetections sensed by each animal. Based on the results of each simulation, FM signals mimicking bat pulses were generated (Fig. 5a–d). The bandwidth and the duration of these signals were fixed at 40 kHz and 3 ms, respectively, and their amplitudes normalized. Therefore, the only difference among signals was their TF. We then calculated cross-correlation functions for a pulse of each bat vis-à-vis the received pulses from other co-flying bats. Signals from other bats were detected when the peaks of the cross-correlation function corresponding to the other bats’ sound exceeded half the maximum value. We then counted the number of detected pulses from others per sensing event. Finally, we considered the single flights in the simulation as group flights without JARs, and conversely the group flights as group flights with JARs.

## Acknowledgments

We thank Hinano Sumiya and Saori Sugihara for their assistance with the data collection in the pilot experiments. We also thank Osama Yamanaka for his assistance with the mathematical modelling. We thank LetPub (www.letpub.com) for linguistic assistance and pre-submission expert review.

Funding: Japan Society for the Promotion of Science KAKENHI grant JP19J02041 (KH), Japan Society for the Promotion of Science KAKENHI grant JP21H05295 (KIK and SH)

## Author Contributions

Conceptualization: KH, SH; Data Curation: KH, TM, NS, SO; Formal analysis: KH; Funding acquisition: KH, SH; Investigation: KH, TM, NS, SO; Methodology: KH, HN, MS; Project administration: SH; Resources: SH; Software: KH, TM, MS, SH; Supervision: SH; Visualization: KH; Writing-original draft: KH; Writing-review & editing: KH, SH, HN, MS, KY, KIK, SH

## Supporting Information for

**Fig. S1.**
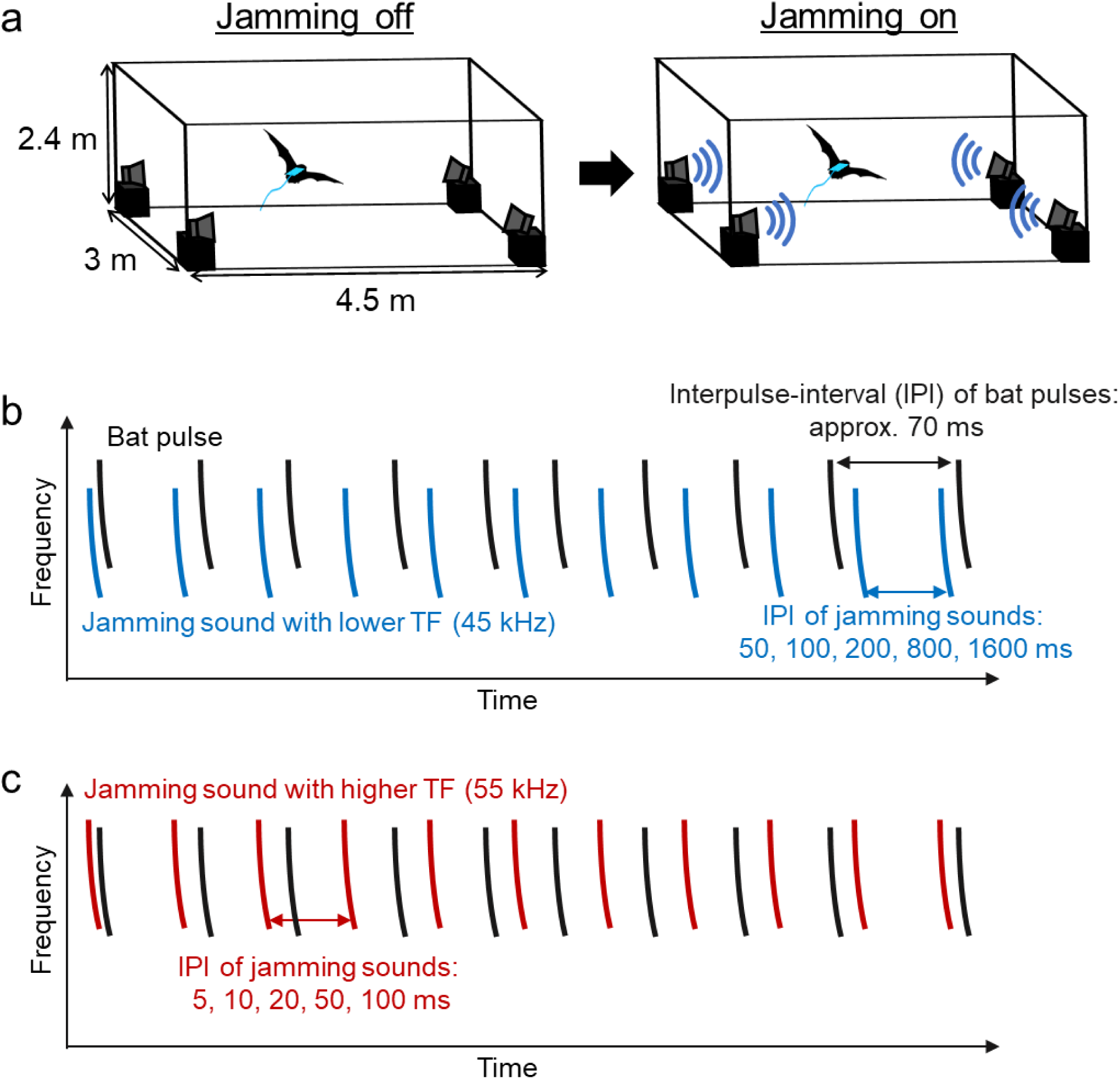
Schematic illustration of the experimental system. (**a**) Experimental setup. Bats carrying a telemetry microphone flew alone in a flight space where four loudspeakers were located at the corners. While flying they were exposed to jamming stimuli presented from the speakers. Schematic spectrograms of jamming stimuli with low TF (**b**, blue) and high TF (**c**, red) in comparison with the bats’ emitted pulses (black).

**Fig. S2.**
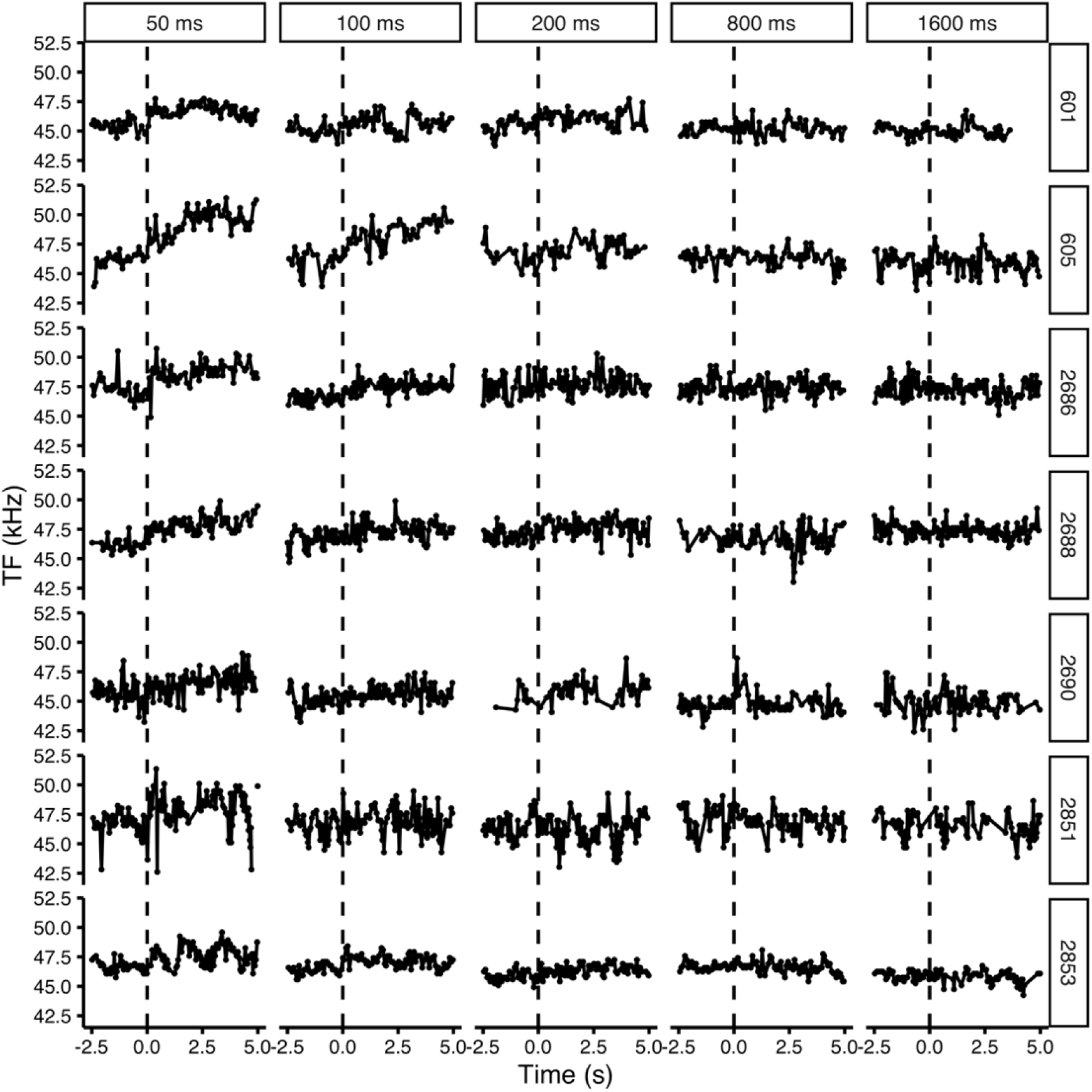
Changes in the TF of individual bats in response to low TF-jamming stimuli with different IPIs. The solid lines indicate the time course of changes in TF of n = 7 bats. The vertical dashed line indicates when the presentation of jamming stimuli started. Likewise for Fig. S3 and S4 below.

**Fig. S3.**
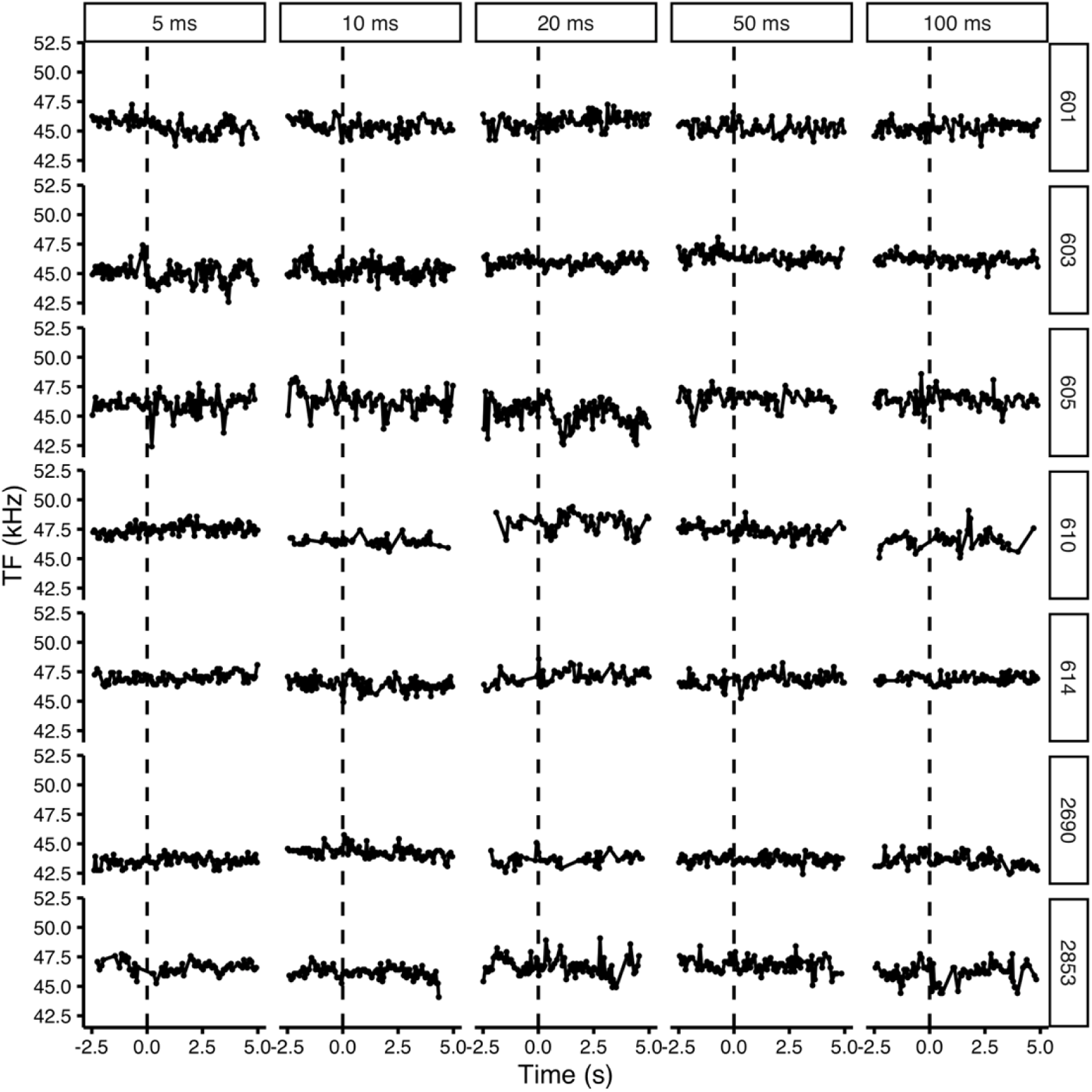
Changes in the TF of individual bats in response to high TF-jamming stimuli with different IPIs.

**Fig. S4.**
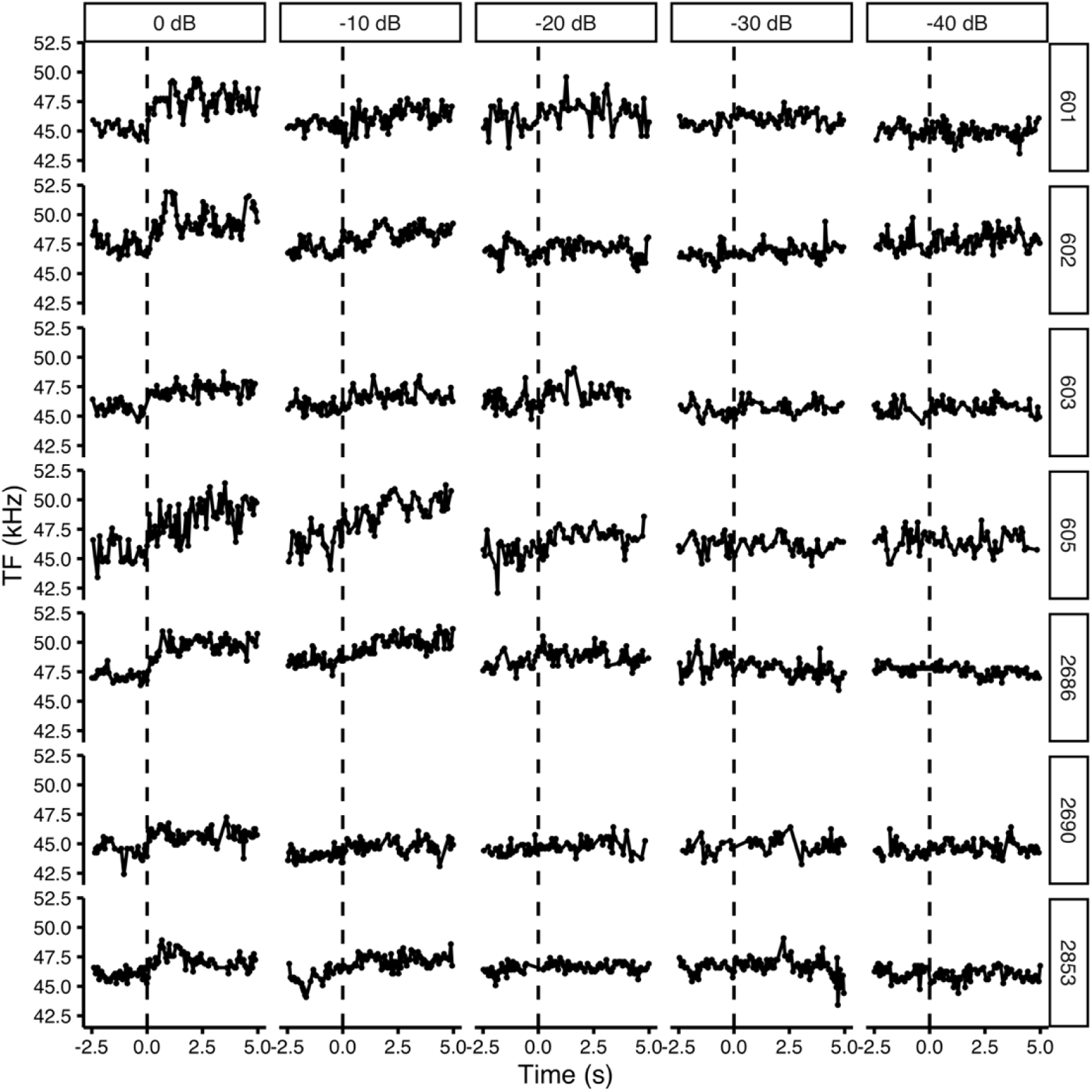
Changes in the TF of individual bats in response to low TF-jamming stimuli with different SPLs.

**Fig. S5.**
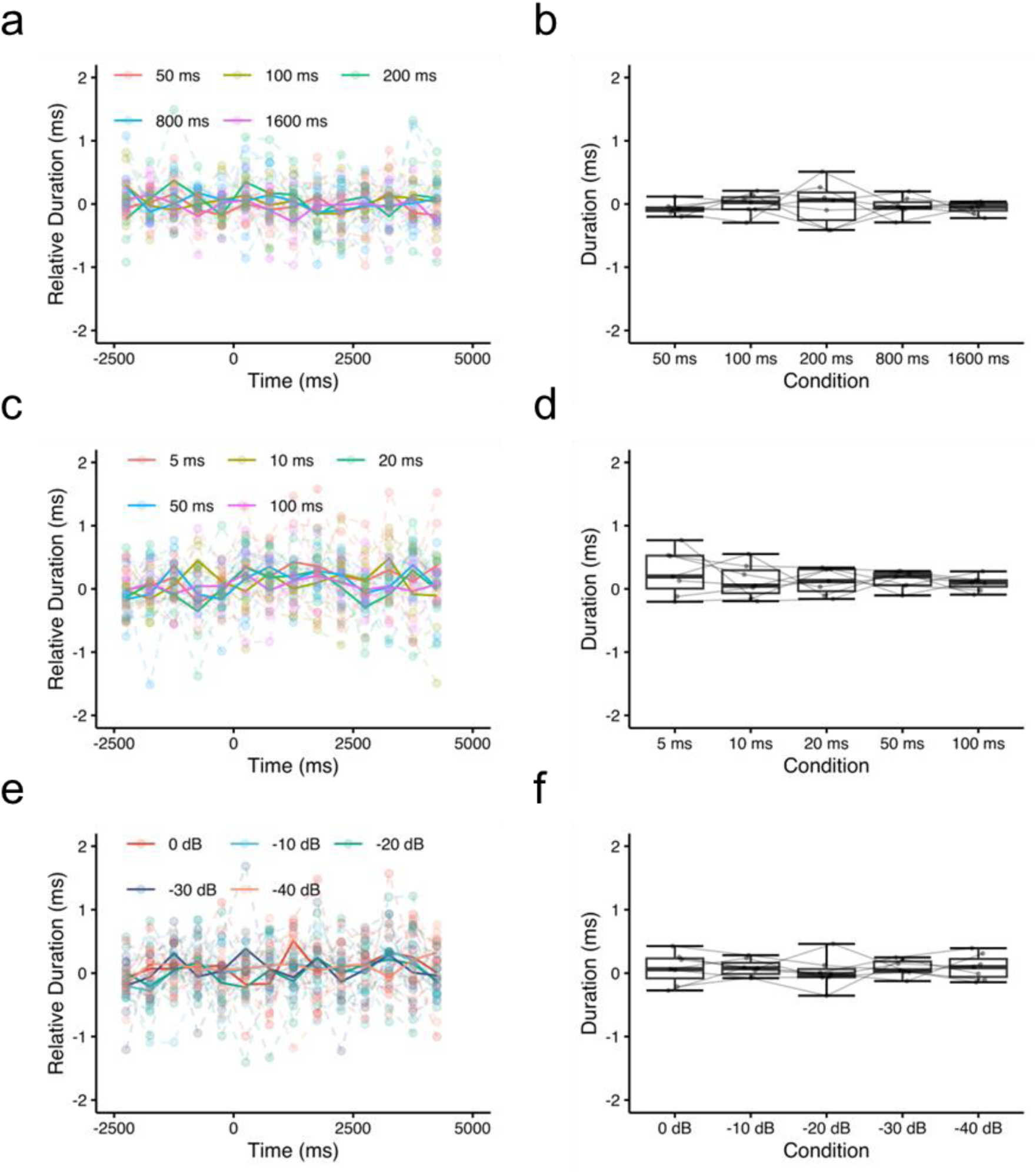
Changes in pulse duration of flying bats in the presence of jamming stimuli. Changes in duration of emitted pulses by bats in response to low TF-jamming stimuli with different IPIs (**a, b**), high TF-jamming stimuli with different IPIs (**c, d**), and low TF-jamming stimuli with different SPLs (**e, f**). Dashed lines and solid lines indicate the individual mean duration and the time course of changes in the population mean of duration, respectively. Colors correspond to different stimulus parameters in **a, c, e**. Plots and lines show mean changes in duration of each individual in **b, d, f**.

**Fig. S6.**
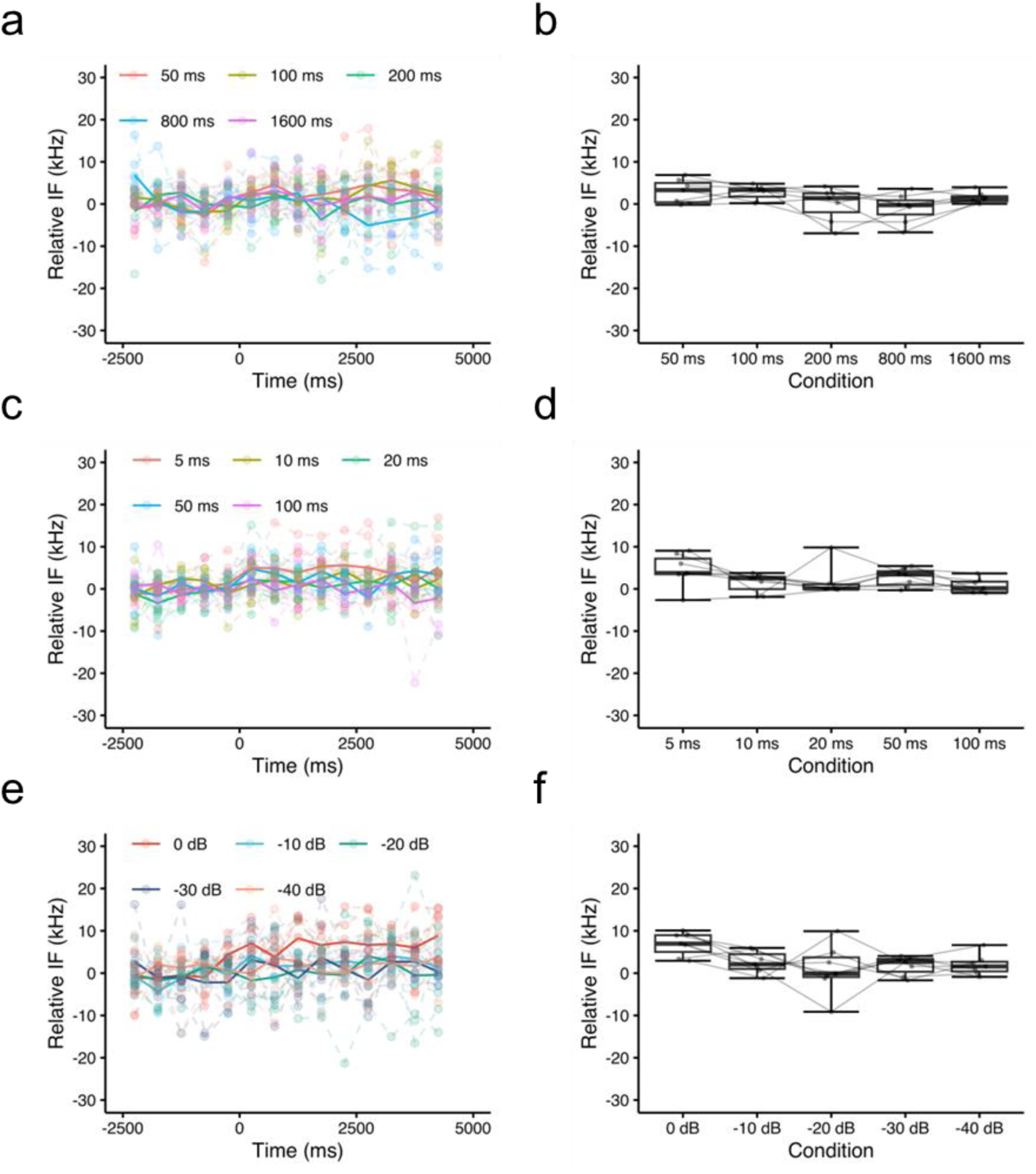
Changes in pulse initial frequency of flying bats in the presence of jamming stimuli. Changes in the initial frequency (IF) of emitted pulses by bats in response to low TF-jamming stimuli with different IPIs (**a, b**), high TF-jamming stimuli with different IPIs (**c, d**), and low TF-jamming stimuli with different SPLs (**e, f**). Dashed lines and solid lines indicate individual mean duration and the time course of changes in the population mean of IF, respectively. Colors correspond different stimulus parameters in **a, c, e**. Plots and lines show mean changes in the IF of each individual in **b, d, f**.

**Fig. S7.**
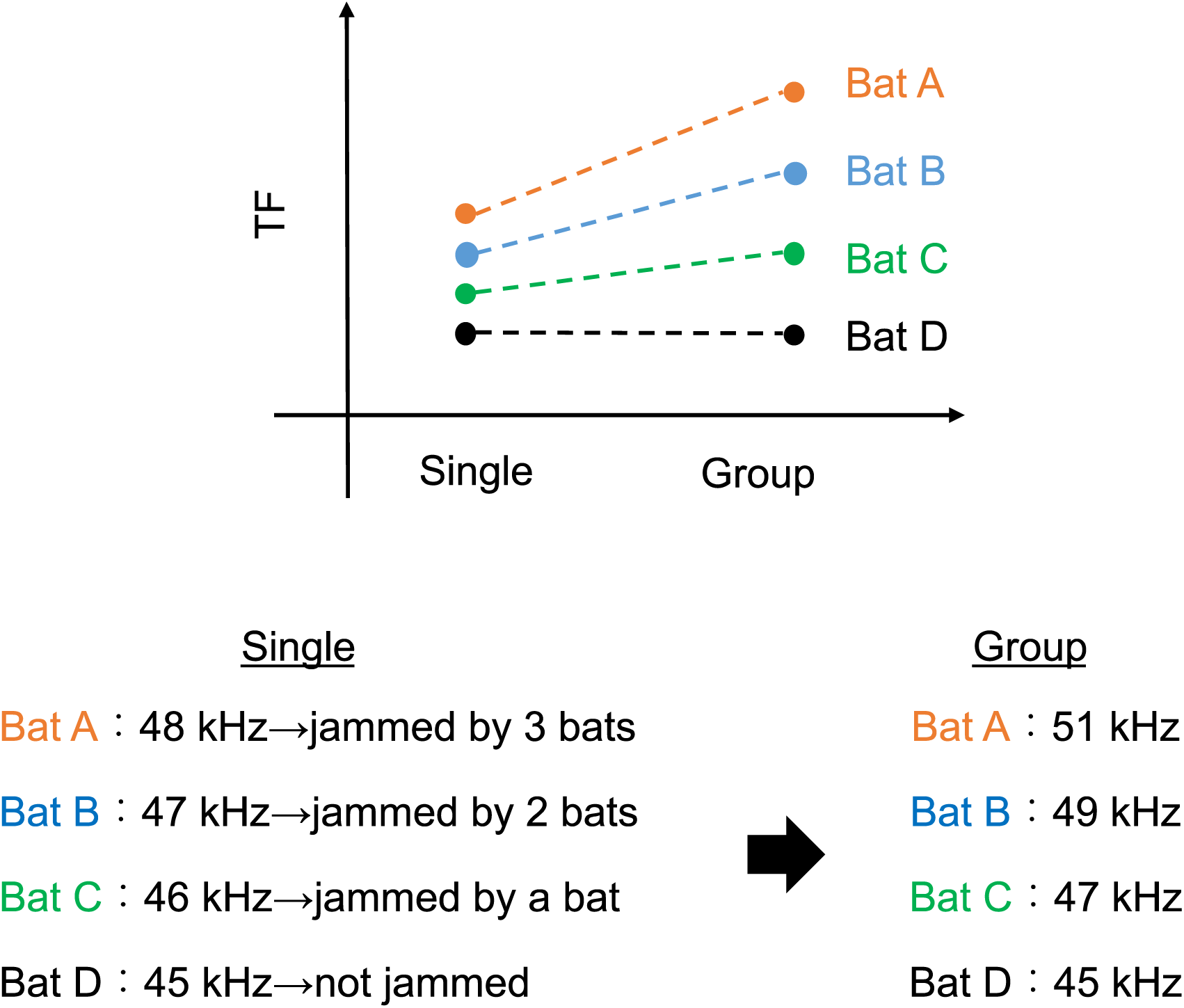
Schematic illustration of how the proposed bat behavioral model works. We assume that bats could be affected by others in flight whose TFs are lower than their own. There are four bats whose TFs varied by 1 kHz. When those four bats flew together, the highest bat (Bat A) is most affected by the other bats and increases its TF the most (3 kHz in this example). The second and third highest-ranked bats increase their respective TF by 2 kHz and 1 kHz; in contrast, the lowest-ranked bat does not change its TF. The resulting mean inter-individual difference thus increases from 1 kHz to 2 kHz without any central command involved.

**Fig. S8.**
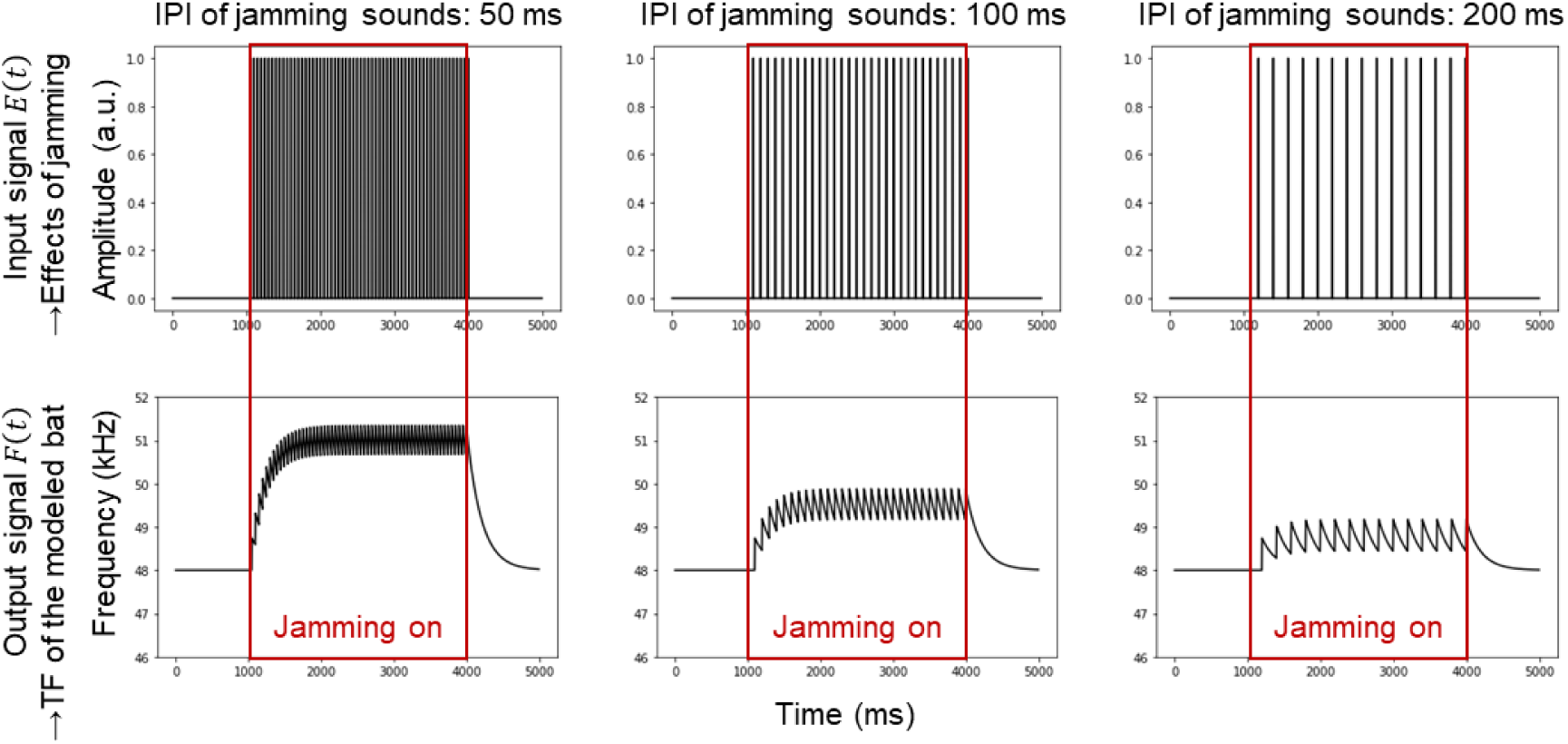
Examples of how the leaky-integrator model works. The received signals are perceived as rectangular pulse trains for bats in the simulations. The model integrates the input with a constant leak, which results in the output frequency increasing more when the IPI of the input signal is higher.

**Fig. S9.**
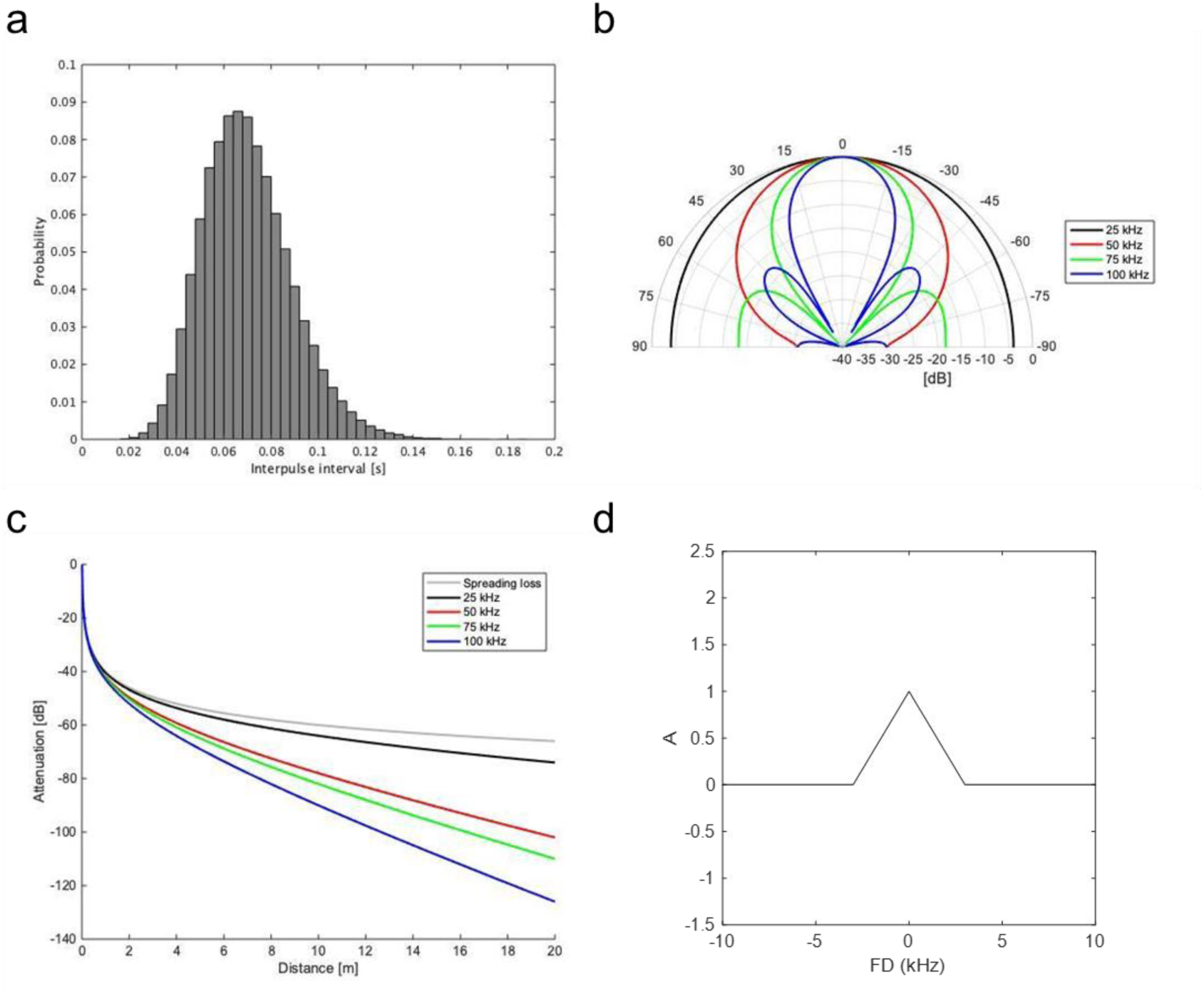
IPIs’ generation, the directional and distance-dependent attenuation, and function in the model simulation. (**a**) The gamma distribution to generate IPIs for the simulations. (**b**) The directionality of pulses in the simulation. For comparison, we used the directionality of 50 kHz for all the emissions. (**c**) The effect of atmospheric attenuation in the simulation. Again, for comparison, we set the attenuation to 50 kHz. (**d**) The shape of the function *A*, which determines how a bat is affected by other bats depending upon frequency difference *FD*.

**Fig. S10.**
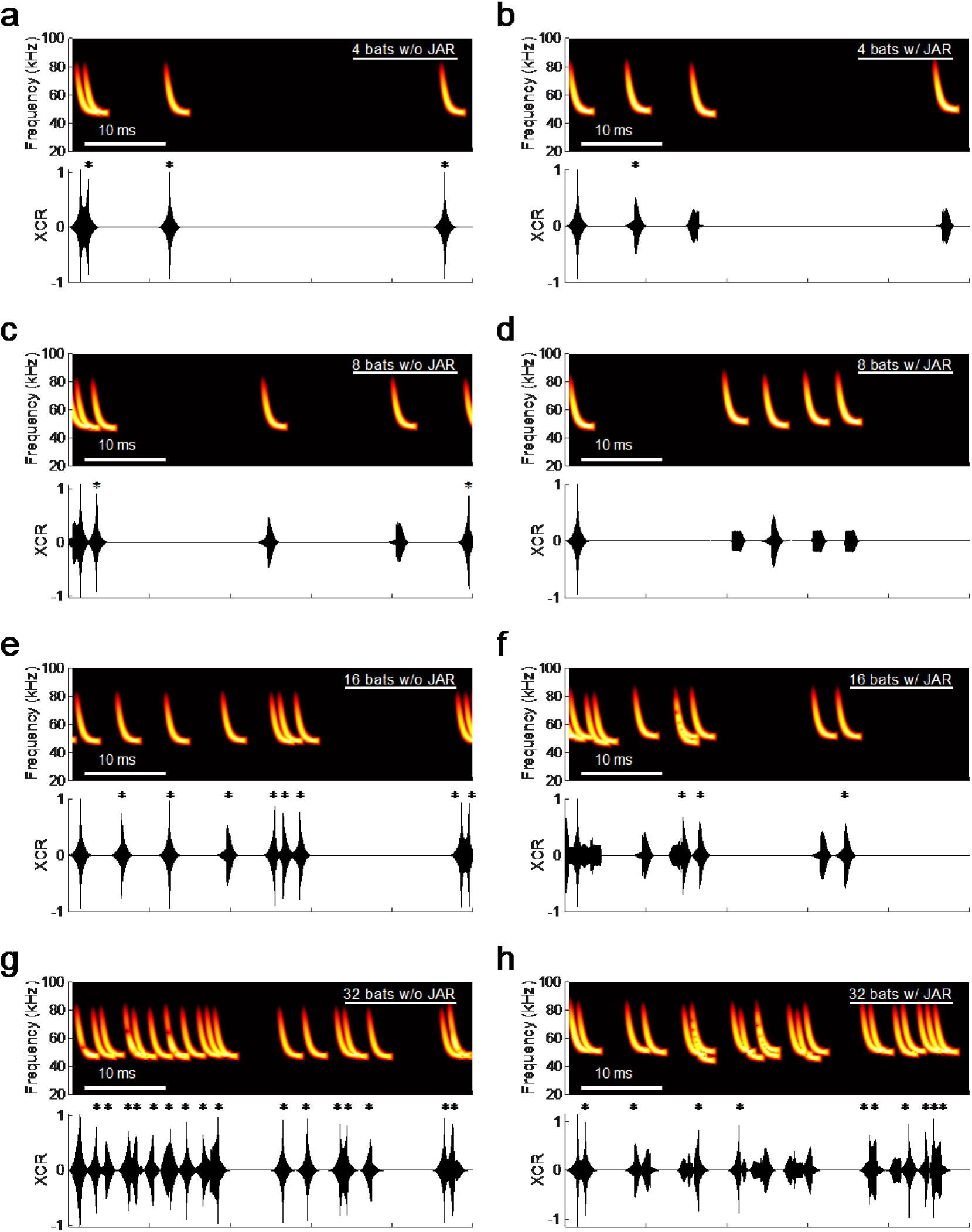
Examples of the effect of JAR on the cross-correlation function. (**a–h**) Example spectrograms (top) and cross-correlation functions (bottom) of a bat’s pulse vis-à-vis other bats’ pulses received by the bat in a group size of 4 (**a**), 8 (**c**), 16 (**e**), and 32 (**g**) bats without JAR (left column) and with JAR (right column). The dashed line in the cross-correlation function represents the threshold for detecting pulses. When pulses from other bats exceed that threshold, the pulses are considered as misdetected (indicated by asterisks). The number of pulses crossing the threshold is smaller in the results with JAR than without JAR employed by bats.

